# Integrative mobilizable elements are pervasive throughout Pseudomonadota

**DOI:** 10.64898/2026.01.12.698999

**Authors:** Adelina Miruna Perta, Francisco Nadal-Molero, Pedro J. Cabello-Yeves, Neris Garcia-Gonzalez, Fernando Gonzalez-Candelas, Alfred Fillol-Salom

## Abstract

Integrative mobilizable elements (IMEs) are mobile genetic elements that reside stably integrated into chromosomes and rely on helper conjugative elements for horizontal transfer. Here, we identify and characterize a widespread family of IMEs, named Pseudomonadota Integrative Mobilizable Elements (PIMEs), which are distributed exclusively in the Pseudomonadota phylum. Genome and phylogenomic analyses reveal ∼1,000 putative PIMEs, comprising at least four distinct PIME subfamilies defined by distinctive genomic organizations and conserved hallmark features. Characterized PIMEs depend on helper conjugative plasmids of the incompatibility group P (IncP) and, upon induction, PIMEs excise, replicate and mobilize intra- and inter-species. Remarkably, the representative PIME and its helper conjugative plasmid engages in cross-complementation, revealing an unrecognized level of functional interplay between hijacker and helper element. We also demonstrate that PIMEs act as reservoirs of known and novel prokaryotic immune systems. Overall, our findings uncover an overlooked and disseminated family of IMEs, which likely plays an important role in bacterial ecology and evolution.

## Introduction

Bacteria are a rich reservoir of a multitude of mobile genetic elements (MGEs), which contribute to the vast intra- and inter-species variations in their gene repertoire^1,2^. These MGEs disseminate from one bacterium to another through a mechanism known as horizontal gene transfer (HGT), a major force in bacterial evolution and pathogenesis, as MGEs frequently carry accessory genes encoding antimicrobial resistances, virulence factors, or prokaryotic immune systems^3–6^.

While some MGEs transfer autonomously between bacterial cells^7–9^, others rely on co-resident MGEs for their mobilization^10–13^. Integrative mobilizable elements (IMEs) represent a discrete class of non-autonomous MGEs that are intimately related to certain conjugative elements, referred to as helper elements, whose life cycle they exploit^10,14^. In the absence of a helper element, IMEs remain stably integrated in the chromosome in a dormant state^15^. Once its helper element is present, IMEs excise from the chromosome, replicate, and hijack the conjugative machinery of their helper elements to mobilize into new bacterial hosts, enabling intra- and inter-species transfer^15–18^.

Our understanding of the IMEs lifecycle is largely derived from a limited number of characterized systems. Indeed, most mechanistic insights into IME-helper interactions have come from *Salmonella* Genomic Island 1 (SGI1) and its derivatives^15,17,19–22^. Members of SGI1 exploit the transcriptional activator AcaCD, encoded by their helper conjugative plasmids of the incompatibility groups A (IncA) and C (IncC), for timing their own excision, replication, and mobilization^10,17,19^. Moreover, SGI1 actively interferes with its helper conjugative plasmids by destabilizing their partitioning systems^23^, by reshaping the conjugative pilus to enhance its own transfer at the expense of the helper^24^, and by interfering with the plasmid relaxase^25^. Whether other IMEs behave similarly or employ other strategies to interact with their helper conjugative elements is yet to be unraveled.

Over the past years, numerous genomic islands have been proposed as putative IMEs based on the presence of an integrase gene combined with the absence of a complete type IV secretion system (T4SS), which is required to mediate conjugative transfer, and/or the presence of a recognizable origin of transfer (*oriT*) region required for mobilization^14,15,20,26^. These putative IMEs comprise diverse families with variable genetic structures and gene repertoires, presumably to adapt to distinct helper conjugative elements^26–29^. However, their helper elements remain unidentified, their complete taxonomic distribution is unknown, the molecular mechanisms underlying their interactions with other MGEs are undetermined, and their impact on their host remains to be discovered.

Here, we report the identification and characterization of a previously unrecognized family of IMEs, Pseudomonadota Integrative Mobilizable Elements (PIMEs), which is widespread across the Pseudomonadota phylum. Members of this family share common genetic modules, and they specifically exploit conjugative plasmids of the incompatibility group P (IncP) as helper elements, which trigger their excision and mediate their intra- and inter-species mobilization. Notably, in contrast to previously described hijacker-helper systems, we show that a helper element co-opts relaxosome functions from an IME to promote its transfer. Furthermore, the accessory regions of these IMEs frequently encode prokaryotic immune systems, suggesting that they confer a potential selective advantage to their hosts. Our study demonstrates that the diversity and distribution of IMEs is underestimated and provide an in-depth characterization of an unrecognized family of IMEs, highlighting that there is still a dearth of fundamental insights governing interactions among MGEs.

## Results

### Identification of novel and widespread IMEs across Pseudomonadota

We identified an uncharacterized element present in *Escherichia coli* strain RHB23-C07. This chromosomally integrated element, with a size of 22.2 kb, encodes genes indicative of mobility: integrase (*int*), transcriptional factors (*alpA* and *luxR-family*), replication-related genes (*repA*), and homologs of conjugative mobilization and transfer genes (*traJ*, *trbJ*, and *trbL*) (Fig.1A). The presence of integration and mobility-associated functions, coupled with the absence of a complete T4SS, suggests that this element may represent either a defective integrative conjugative element (ICE) or an integrative mobilizable element (IME)^14,30^. To investigate this, we assessed the similarity between this element and the genomes of typical ICEs and IMEs retrieved from ICEberg^27^, a comprehensive database of both experimentally validated and predicted MGEs. Our analysis shows that the element found in *E. coli* RHB23-C07 is different from other known ICEs and IMEs (Table S1). While the resemblance to ICEs is limited to those that encode *int* and *alpA* as core genes, or to misannotated ICEs that are, in fact, IMEs (see Table S1), this element shows partial resemblance to some predicted IMEs: GIsul2, BcenGI2, and IncP island^31–34^, particularly in the *int* and *alpA* genes, likely due to some of these MGEs integrating at the same *attB* site. Yet, the overall sequence similarity is low and restricted to genes indicative of mobility (Fig. 1A, Table S1). Given that these predicted IMEs differ in gene content and organization, we hypothesized that they could form part of a broader IME family comprising distinct subfamilies, and the element found in *E. coli* RHB23-C07 may therefore represent a previously uncharacterized subfamily within these IMEs.

**Figure 1.**
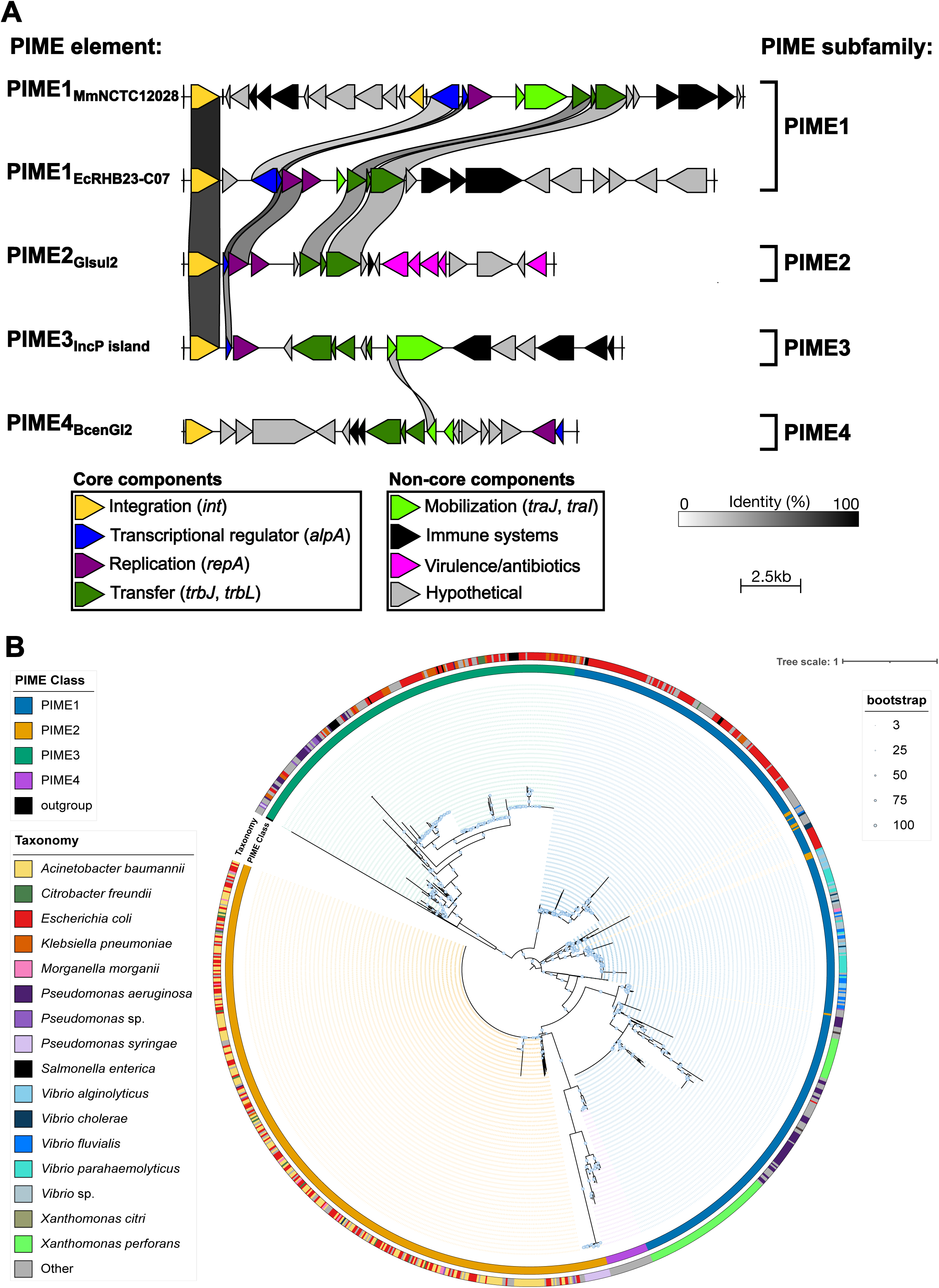
Distribution and gene content of PIME-like elements. **(A)** Schematic representation of representative PIME elements. Genes are color-coded according to their sequence and predicted function: integrase (*int*) in yellow; transcriptional regulators (*alpA* and *luxR*) in blue; replication genes (*repA* and *repC*) in purple; mobilization genes (*traK*, *traJ,* and *traI*) in light green; transfer genes (*traC*, *trbJ*, *trbK,* and *trbL*) in dark green; virulence/antibiotic resistance in pink; immune systems in black; and genes encoding hypothetical proteins in grey. Grey shading between PIME elements indicates regions of shared sequence similarity. Comparisons and visualizations were generated using Clinker. **(B)** A phylogenetic core tree representing 1136 putative PIME elements found in bacterial genomes from the RefSeq database. Colours on the outer ring represent the species where the PIME element is present and the PIME subfamily.

To characterize the distribution and diversity of this IME family, we assembled a representative collection of putative IMEs by performing genome searching using the conserved core components shared by the distinct subfamilies identified (Fig. 1A). Specifically, we searched for the co-occurrence of an integrase, *alpA*, a replicase, and conjugative genes (*trbJ* and *trbL*) across the RefSeq database of bacterial genomes (see methods for more details). We identified a total of 1136 elements that contain all core components within genomic regions from almost 100 Pseudomonadota species (Fig. 1B, Table S2). Most of these putative IMEs (94.05%) were detected in Gammaproteobacteria, predominantly within the Enterobacteriaceae family, including *E. coli*, *Klebsiella pneumoniae*, *Morganella morganii*, or *Enterobacter hormaechei*; the Vibrionaceae family, such as *Vibrio alginolyticus*, *Vibrio cholerae, Vibrio parahaemolyticus, or Vibrio vulnificus*; the Pseudomonadaceae family, including *Pseudomonas aeruginosa* and *Pseudomonas syringae*; and the Moraxellaceae family, including *Acinetobacter baumanii*. This is due to the fact that more than 42% of all the bacterial genomes in the dataset belong to the Gammaproteobacteria

class. Additional putative IMEs were identified in other members of the Pseudomonadota phylum such as Betaproteobacteria. Notably, the different IME subfamilies do not tend to associate with specific bacterial hosts, suggesting extensive horizontal gene transfer across phylogenetically distant bacteria. In addition, the lack of clustering among subfamilies indicates a low frequency of recombination events between distinct IME subfamilies (Fig. 1B, Table S2).

Given that this IME family is found exclusively within Pseudomonadota, we designated it as PIME (<u>P</u>seudomonadota Integrative <u>M</u>obilizable <u>E</u>lement). Individual PIME elements are named with reference to their subfamily (see methods for more details), species and specific strain. Thus, the element identified in *E. coli* strain RHB23-C07 would be referred to as PIME1EcRHB23-C07 (<u>P</u>seudomonadota Integrative <u>M</u>obilizable <u>E</u>lement Subfamily <u>1</u> *<u>E</u>scherichia <u>c</u>oli* strain RHB23-C07). Despite their diversity, these putative PIMEs share several defining features: (i) modularity, with common genetic modules related to integration, regulation, replication, and transfer; (ii) mobilization strategies, which vary across members, as some encode *traJ* or a relaxase (*traI*), while others do not; (iii) distinct subfamily signatures, with each subfamily displaying a characteristic genetic organization, as exemplified by PIME3 and PIME4, or conserved hallmark features, as observed for PIME1, which encodes two divergently oriented transcriptional regulators within its regulatory region. One regulator corresponds to *alp*A, whereas the other is a member of the *luxR* family of transcriptional regulators; (iv) large accessory regions, which can comprise over 50% of the element’s genome and frequently carry genes associated with prokaryotic immune systems; and (v) prevalence across species, with identical or closely related putative PIMEs found in multiple bacterial species, suggesting mobility between species and genera. For example, PIME1EcRHB23-C07 is found in multiple *E. coli* genomes, as well as in *Citrobacter meridianamericanus* and *Shigella sonnei* (Fig. S1). Similarly, a related element from *M. morganii* strain NCTC12028, referred to as PIME1MmNCTC12028, is found in *Providencia rettgeri*, *Providencia huaxiensis*, and *Proteus vulgaris* (Fig. 1A, Fig. S1).

### Excision of PIMEs depends on helper conjugative plasmids

Given the widespread distribution of these IMEs, we sought to characterize their biology. To investigate whether putative PIMEs, integrated adjacent to the *guaA* gene, undergo spontaneous excision and circularization, we selected two representative elements from the novel subfamily found across multiple species, PIME1EcRHB23-C07 and PIME1MmNCTC12028 (Fig. S1). The main difference between them is that PIME1MmNCTC12028 encodes a relaxase (*traI*) whereas PIME1EcRHB23-C07 does not (Fig. 1A). We performed PCR analysis using inward- and outward-directed primers (Fig. S2) and we did not observe spontaneous excision and circularization. This result suggests that both elements either lack excision ability or require additional external factors to facilitate excision.

Next, we tested whether co-resident conjugative plasmids could trigger the excision of these putative PIMEs. We introduced various conjugative plasmids belonging to multiple incompatibilities groups into *E. coli* RHB23-C07 and *M. morganii* NCTC12028 strains (Table S3). Then, we performed PCR analyses to assess integration, circularization and excision, as previously performed. Notably, pB10 and pKJK5 conjugative plasmids successfully promoted the excision and circularization of PIME1EcRHB23-C07 and PIME1MmNCTC12028, whereas RIP113, pCF12 and pOXA-48 plasmids did not (Fig. 2A and Fig. S2D). This indicates that pB10 and pKJK5 could function as helper conjugative plasmids, promoting the PIME excision and circularization. Interestingly, both conjugative plasmids belong to the IncP group, whereas RIP113 is classified as IncN, pCF12 as IncX, and pOXA-48 as IncL/M, suggesting that the ability to facilitate excision may be specific to the IncP group plasmids.

**Figure 2.**
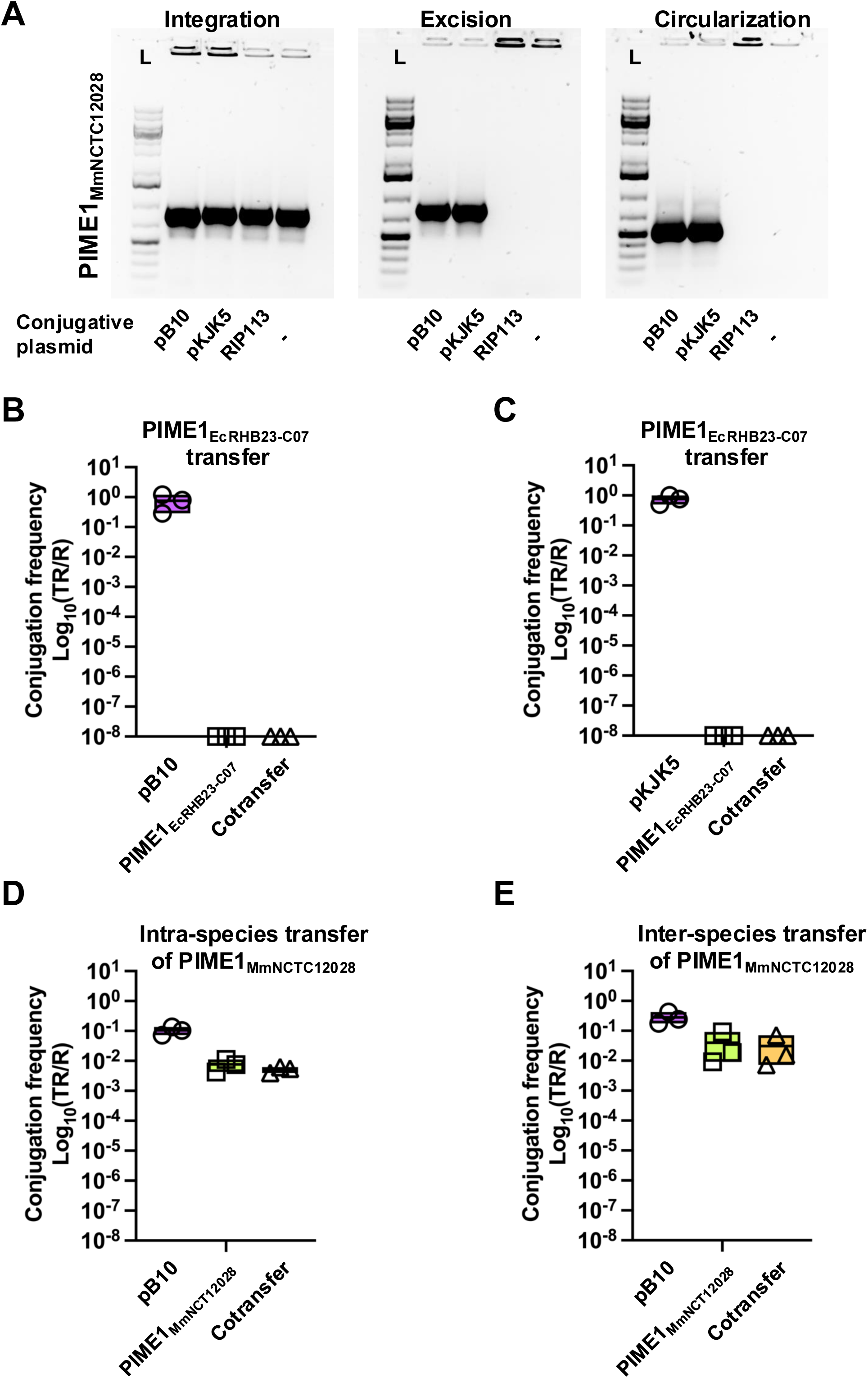
Intra- and inter-species mobilization of PIME1_MmNCTC12028_. **(A)** Genomic DNA from *M. morganii* NCTC12028 carrying different conjugative plasmids (pB10, pKJK5, and RIP113) was extracted and PCR-amplified using specific primers designed to detect integration (external and internal sequences of the PIME element), excision (chromosomal flanking regions), and circularization (pair of divergently oriented primers at the PIME termini). L, DNA ladder (AMPIGENE, 250-10,000bp). **(B, C)** Mobilization of PIME1_EcRHB23-C07_ to *E. coli* c1a. Conjugation assays were performed with PIME1_EcRHB23-C07::*cat*_ carrying helper conjugative plasmids pB10 (B) or pKJK5 (C) and the recipient strain *E. coli* c1a (*yciO::kmR*). Transfer of conjugative plasmid, PIME1_EcRHB23-C07_, and their cotransfer was evaluated. Values represent the transconjugants per recipient cells (TR/R). Error bars indicate standard deviation (*n = 3*). **(D, E)** Mobilization of PIME1_MmNCTC12028_ to different species. Conjugation assays were performed with PIME1_MmNCTC12028::*tetA*_ carrying a derivative helper conjugative plasmid pB10::*cat* and the recipient strain *M. morganii* NCTC12028 ΔPIME (D) or *E. coli* c1a (*yciO::kmR*). Transfer of pB10, PIME1_MmNCTC12028_, and their cotransfer was evaluated. Values represent the transconjugants per recipient cells (TR/R). Error bars indicate standard deviation (*n = 3*).

### Intra- and inter-species transfer of PIMEs

The previous result suggests that conjugative plasmids pB10 and pKJK5 could act as helper elements for both PIMEs. To assess the mobilization of these PIMEs *in vivo*, we inserted antibiotic resistance markers into each element: a *cat* selection marker into PIME1EcRHB23-C07, and a *tetA* selection marker into PIME1MmNCTC12028. The localization of the antibiotic resistance markers is at the 3’ end of their genomes (Fig. S3A). For PIME1EcRHB23-C07 element, we analyzed whether the PIME could be mobilized in the presence of its helper plasmids, pB10 and pKJK5, both of which naturally encode a *tetA* marker. Surprisingly, while helper plasmids themselves were highly transferred, the PIME was not mobilized (Fig. 2B and 2C). Note that similar results were observed regardless of whether the recipient strain was *E. coli* c1a (a lab adapted strain) or *E. coli* RHB23-C07 with scarless deletion of the PIME to minimize potential HGT barriers in recipient strains (Fig. S3B and S3C). This indicates that both helper conjugative plasmids can promote PIME1EcRHB23-C07 element induction but not their mobilization, suggesting that it likely depends on a compatible helper conjugative plasmid or an additional MGE for mobilization.

Next, we analyzed whether PIME1MmNCTC12028 is mobilized in the presence of a derivative pB10 helper plasmid, in which the native *tetA* marker was replaced with a *cat* marker. Using a recipient strain of *M. morganii* NCTC12028 with a scarless deletion of the PIME, we observed that both MGEs, pB10 and PIME1MmNCTC12028, were highly mobilized (Fig. 2D). Given that the PIME1MmNCTC12028 element is found in multiple bacterial species, to examine inter-species transfer, we used *E. coli* c1a as the recipient strain and we obtained similar transconjugation frequencies (Fig. 2E). Integration of PIME1MmNCTC12028 into the *guaA* locus of *E. coli* was confirmed by PCR (Fig. S3D).

### Characterization of the genetic modules in PIME1MmNCTC12028

Although both PIMEs tested are induced by helper pB10 conjugative plasmid, only the PIME1MmNCTC12028 element is mobilized at high frequencies. We wanted to further characterize the PIME1MmNCTC12028-encoded genes to determine the role of the main distinct genetic modules in the PIME’s lifecycle: (i) integration, (ii) excision, (iii) replication, and (iv) conjugative mobilization. To test this, we generated individual mutations in genes associated to integration, regulation, replication, and mobilization (Fig. 3A). We then assessed the ability of each PIME1MmNCTC12028 mutant element, in the presence of a coresident pB10 *cat* helper conjugative plasmid, to transfer to new hosts through conjugation experiments. The transconjugation efficiency of PIME1MmNCTC12028 was affected in all mutants of its main genetic modules tested.

**Figure 3.**
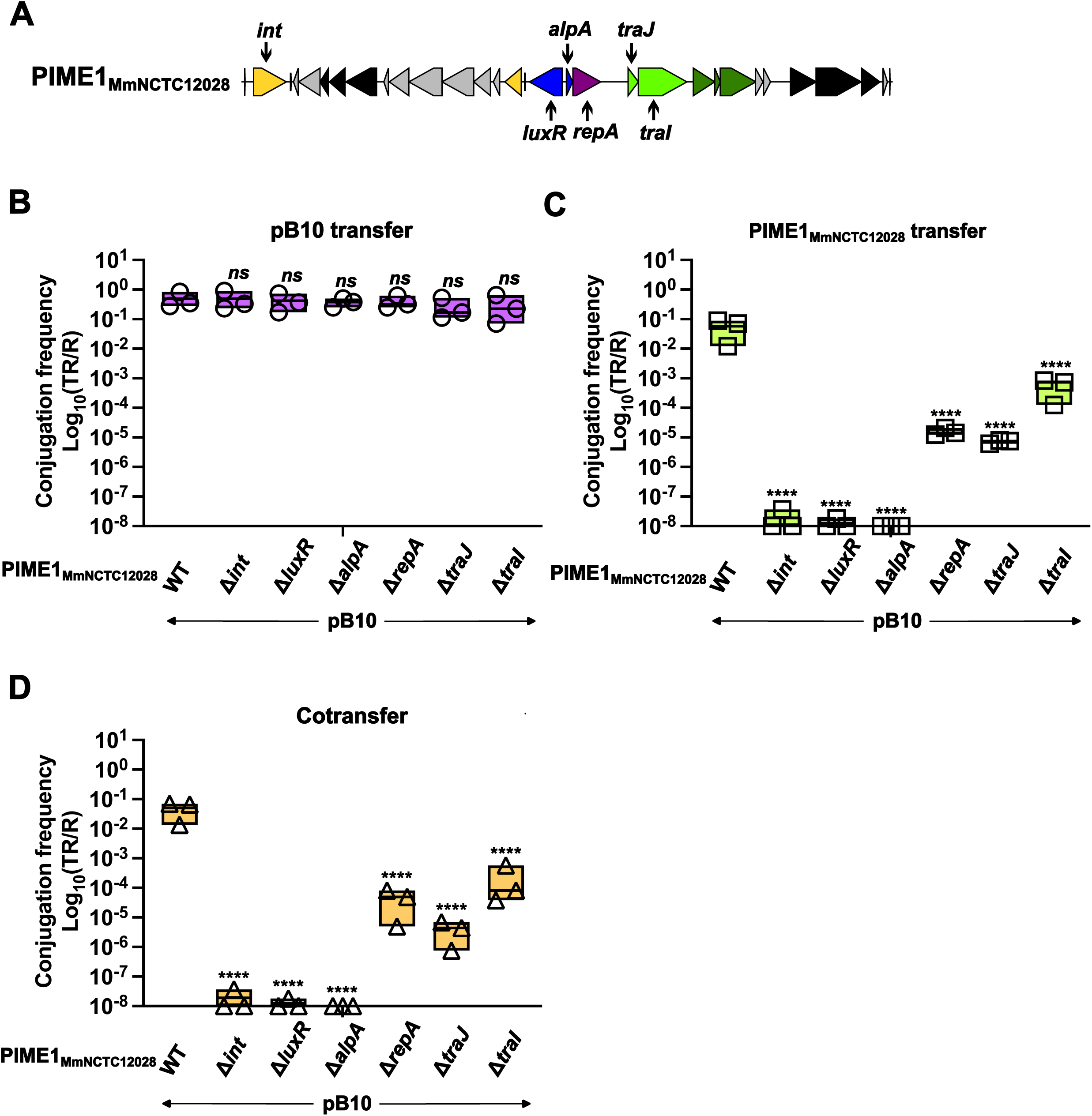
Effect of different IME gene mutations on pB10, PIME1_MmNCTC12028_, and cotransfer. **(A)** Genetic map of PIME1_MmNCTC12028_, with arrows indicating the genes mutated. **(B-D)** Mobilization of PIME1_MmNCTC12028_. Conjugation assays were performed with different PIME1_MmNCTC12028::*tetA*_ derivatives carrying mutations in *int*, *luxR*, *alpA*, *repA*, *traJ*, and *traI*, and in presence of the helper conjugative plasmid pB10::*cat*. *E. coli* c1a (*yciO*::*kmR*) was used as the recipient strain. Transfer of pB10 (B), PIME1_MmNCTC12028_ (C), and their cotransfer (D) was evaluated. Values represent the transconjugants per recipient cells (TR/R). Error bars indicate standard deviation (*n* = *3*). For (B-D), data were log_10_ transformed prior to analysis. A one-way ANOVA with Dunnett’s multiple comparisons test was performed to compare mean differences between WT and individual mutants. Adjusted *p* values were as follows: *ns* > 0.05; \**p* ≤ 0.05; \*\**p* ≤ 0.01; \*\*\**p* ≤ 0.001; \*\*\*\**p* ≤ 0.0001.

While mutations in the replicase and mobilization genes severely affected its transfer, mutants in the integrase and regulatory genes completely abolished it (Fig. 3C). To verify that these phenotypes were consequence of the gene deleted, each mutant was complemented *in trans* with the corresponding gene cloned in the expression vector pBAD18, under control of the arabinose-inducible promoter (*PBAD*). In all cases, conjugative transfer of the PIME1MmNCTC12028 element was partially restored (Fig. S4).

Since deletion of *int*, *luxR*, and *alpA* completely abolished PIME1MmNCTC12028 transfer, we next analyzed the ability of those mutants to excise and circularize, as determined by qPCR analysis. Neither excision nor circularization was detected in any of the mutants (Fig. S5), suggesting that these genes are essential for these processes. To confirm the previous results, we collected the total chromosomal DNA from individual mutants in integration, regulation, and replication in the presence of a coresident pB10 plasmid, and subjected them to whole-genome sequencing. We quantified the reads corresponding to chromosome, PIME1MmNCTC12028, and pB10 and represented them as depth of coverage relative to the average of the entire genome (Fig. 4). While PIME1MmNCTC12028 showed strong amplification up to eight times compared to the chromosome in the presence of the helper pB10 conjugative plasmid, amplification of PIME1MmNCTC12028 Δ*repA* was highly affected. In contrast, PIME1MmNCTC12028 Δ*int*, Δ*luxR*, and Δ*alpA* were incapable to replicate (Fig. 4). These results are consistent with the qPCR and transconjugation data (Fig. 3 and Fig. S5). Although integrase is known to mediate excision and could explain the lack of amplification as the PIME is not able to replicate extrachromosomally, it is tempting to speculate that both transcriptional regulators may play a role in recognizing the presence of helper conjugative plasmids. The identification of the helper plasmid inducer and the underlying mechanism is currently under investigation.

**Figure 4.**
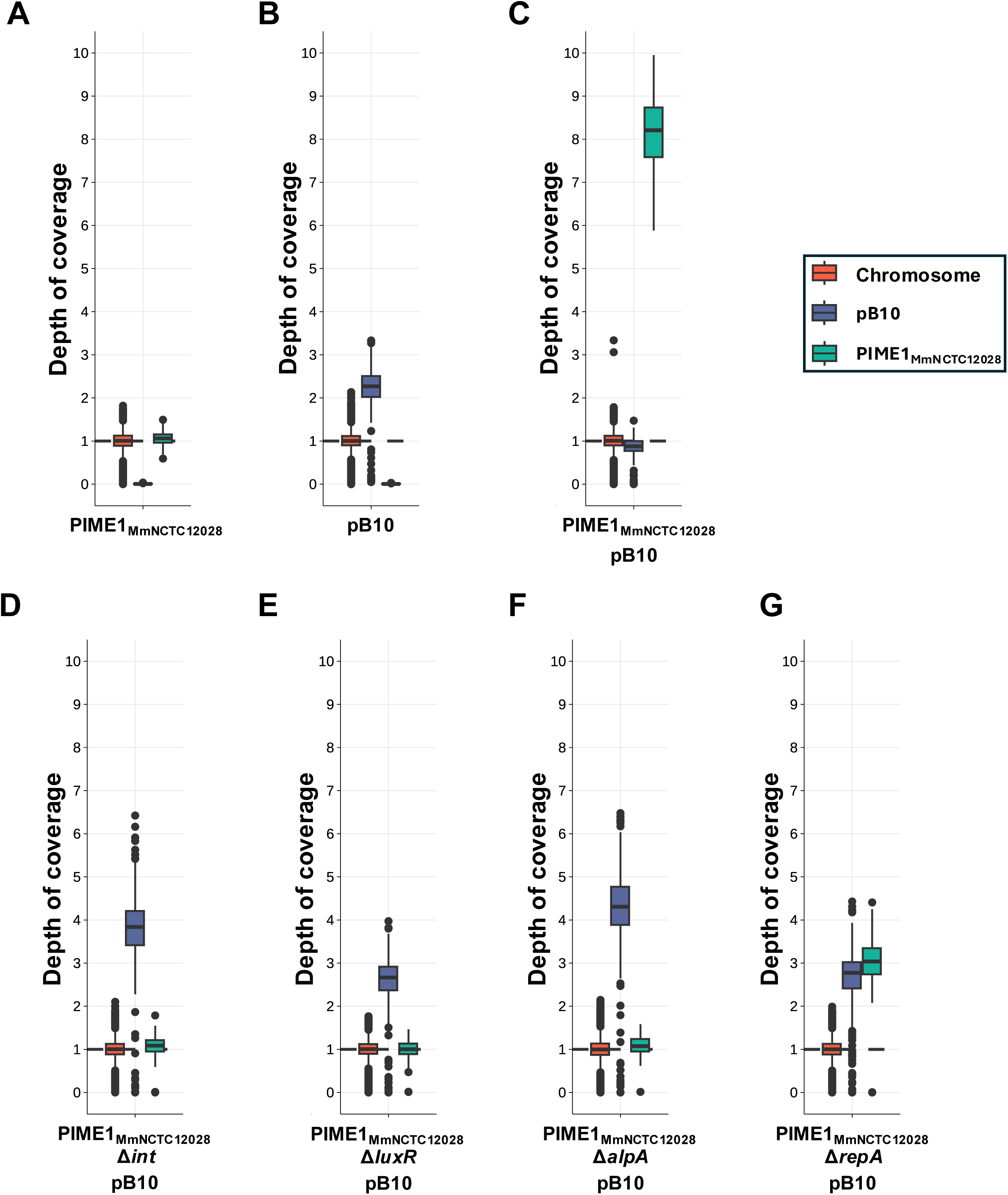
Monitoring PIME1_MmNCTC12028_ and pB10 abundance by whole genome sequencing. Relative depth of coverage of the bacterial chromosome, PIME1_MmNCTC12028_, and the conjugative plasmid pB10. Total genomic DNA was collected and sequenced from strains carrying WT PIME1_MmNCTC12028_ alone (A), pB10 alone (B), WT PIME1_MmNCTC12028_ in the presence of pB10 (C), or individual PIME1_MmNCTC12028_ mutants in integration (*int*) (D), regulation *luxR* (E), *alpA* (F), or replication (*repA*) (G) in the presence of a coresident pB10 plasmid. Relative depth of coverage represents DNA abundance normalized to the average bacterial genomic depth of coverage.

Notably, unlike SGI1^15,35^, we did not observe any reduction in the cotransfer of the pB10 conjugative plasmid and the PIME1MmNCTC12028 element (Fig. 2, Fig. 3 and Fig. S4). However, the presence of PIME1MmNCTC12028 affected pB10 amplification. Specifically, we observed that pB10 amplification was dramatically reduced in the presence of PIME1MmNCTC12028 wild-type, but not its mutants in the integrase, regulatory, and replicase genes (Fig. 4). These results indicate that the reduction in helper conjugative plasmid amplification mediated by PIME1MmNCTC12028 requires both excision and active replication of the PIME element. Furthermore, colony size was reduced in strains harboring both elements only when PIME1MmNCTC12028 was capable to excise and replicate (Fig. S6). Together, these observations suggest a potential incompatibility between the two elements, despite the absence of any detectable effect on conjugative plasmid transfer efficiency (Fig. 3A).

### Identification of the origin of replication (*oriV*) of PIME1MmNCTC12028

The previous results suggested that PIME1MmNCTC12028 encodes a functional replicon and its replication is impaired in the absence of its cognate replicase (PIME1MmNCTC12028 Δ*repA*; Fig. 3 and 4). To identify the replicon organization, we examined intergenic regions and identified a putative *oriV* locus immediately downstream of *repA* (Fig. S7A). This *oriV* locus resembles the origin of typical iteron-containing replicons: containing four direct repeats (iterons) and a putative DnaA box flanked by an AT-rich region (75%). A second AT-rich region (60%) flanking the iterons contains two inverted repeats and one direct repeat. To demonstrate the *repA-oriV* essentiality for PIME replication, we cloned the *repA-oriV* regions from PIME1MmNCTC12028 into the thermosensitive pK03-Blue plasmid, under the control of an arabinose-inducible promoter. Note that this plasmid cannot grow at restrictive (44 °C) temperature. We also constructed plasmids carrying mutations in either the *repA* gene or the *oriV* locus. We tested the ability of these plasmids to grow at the permissive (30 °C) and the restrictive (44 °C) temperatures. Only strains carrying the intact *repA-oriV* region generated colonies at the restrictive temperature (Fig. S7B). Note that at restrictive temperatures, the number of colonies obtained was reduced compared to permissive temperatures, suggesting defects in plasmid segregation. These results confirm that PIME1MmNCTC12028 encodes a functional replication module and that the sequence downstream of *repA* contains the *oriV* locus.

### PIMEs contain an *oriT* region similar to their helper conjugative plasmids

Given that both PIMEs analyzed are induced by helper conjugative plasmids of the IncP group, we examined their genomes to determine whether they carry an *oriT* region similar to that of their helper conjugative plasmids. IncP plasmids possess several essential features within their *oriT* regions: (i) symmetric 19 bp sequence repeats; and (ii) a *nic* region containing a conserved *nic* site^36,37^. We observed that both elements, PIME1EcRHB23-C07 and PIME1MmNCTC12028, carry an *oriT* region resembling that of IncP plasmids (Fig. S8). Although the *nic* site is conserved, both PIMEs contain unique inverted repeats of different size adjacent to the *nic* site: 17 bp in PIME1EcRHB23-C07 and 20 bp in PIME1MmNCTC12028. Notably, these inverted repeats are recognized by TraJ, which binds to these sequences and forms a nucleoprotein structure that initiates the assembly of a functional relaxosome^38^. The presence of *traJ* genes in both PIME genomes may explain these unique inverted repeats in PIMEs (Fig. 1 and Fig. S8).

Although PIME1EcRHB23-C07 is induced by helper conjugative plasmids, it is not mobilized by them. Thus, it is tempting to speculate that this lack of mobilization could result from either: (i) incompatibility between the PIME-encoded TraJ and the conjugative plasmid relaxosome, or (ii) the plasmid relaxosome does not recognize the PIME-specific inverted repeats. To test this, we generated a series of mobilizable plasmids by independently cloning the *oriT* regions from pB10, pKJK5, PIME1EcRHB23-C07, and PIME1MmNCTC12028 into the non-transmissible plasmid pBAD18-*kmR*, which cannot be mobilized by pB10 and pKJK5 (Fig. S9). Note that the *traJ* gene from PIME1EcRHB23-C07, as well as the *traJ* and *traI* genes from PIME1MmNCTC12028, were cloned within their respective *oriT* regions, under the control of an arabinose-inducible promoter. Each construct was then introduced into *E. coli* strains carrying either pB10 or pKJK5, and conjugation assays were performed to measure the transconjugation frequencies. The conjugation frequency of pB10 itself was unaffected regardless the mobilizable plasmid version that carried it. Among the mobilizable plasmids, we observed successful mobilization for those versions carrying the *oriT* regions of pB10 and PIME1MmNCTC12028. In contrast, low mobilization was observed for the PIME1EcRHB23-C07 *oriT* region (Fig. S9A). Similar results were obtained using plasmid pKJK5 (Fig. S9B). These findings indicate that the relaxosome of IncP helper conjugative plasmids does not recognize the PIME1EcRHB23-C07 *oriT* region and that PIME1EcRHB23-C07-TraJ is unable to hijack the helper plasmid relaxosome, suggesting that PIME1EcRHB23-C07 likely relies on a distinct helper conjugative plasmid relaxosome for mobilization.

Finally, we confirmed that PIME1MmNCTC12028 relies on its own relaxase for mobilization, as deletion of the PIME *traI* gene significantly reduced the mobilizable plasmid transfer (Fig. S9). We next assessed the transfer efficiency of the PIME1MmNCTC12028 mobilizable plasmid in a pKJK5 Δ*traI* strain. We hypothesized that, in this condition, the plasmid containing the PIME *oriT* would remain efficiently mobilized, while transfer of pKJK5 Δ*traI* would be impaired. Unexpectedly, although the PIME1MmNCTC12028 *oriT* plasmid was efficiently mobilized, we also observed pKJK5 Δ*traI* transconjugants, but only in the presence of PIME1MmNCTC12028 TraI (Fig. S9C). Collectively, these results suggest that PIME1MmNCTC12028 depends on its own relaxase for mobilization, but pKJK5 can also exploit the relaxase from PIME1MmNCTC12028 to mediate its own transfer.

### PIMEs are reservoirs of diverse prokaryotic immune systems

One notable feature of these PIMEs is the size of their accessory regions, which comprise over 50% of their genomes and it is generally located adjacent to the transfer module (Fig. 1). Using DefenseFinder^39^, we analyzed the accessory regions of the characterized PIME1EcRHB23-C07 and PIME1MmNCTC12028 elements and we identified multiple prokaryotic immune systems (Fig. 5A). While PIME1EcRHB23-C07 encodes a Lamassu system^40^, PIME1MmNCTC12028 encodes a retron system (retron-Eco7)^41^. In addition, PIME1MmNCTC12028 contains another accessory region located between the integrase and the regulatory modules and it contains a restriction modification system. Given that PIMEs are mobilized between species, we first tested whether PIME1MmNCTC12028 confers protection to its new *E. coli* host against phage predation. However, we did not observe protection against any of the phages tested (Fig. 5B). Because this PIME is not naturally found in *E. coli*, it is plausible that the encoded immune systems are not adapted to block phages in *E. coli*.

**Figure 5.**
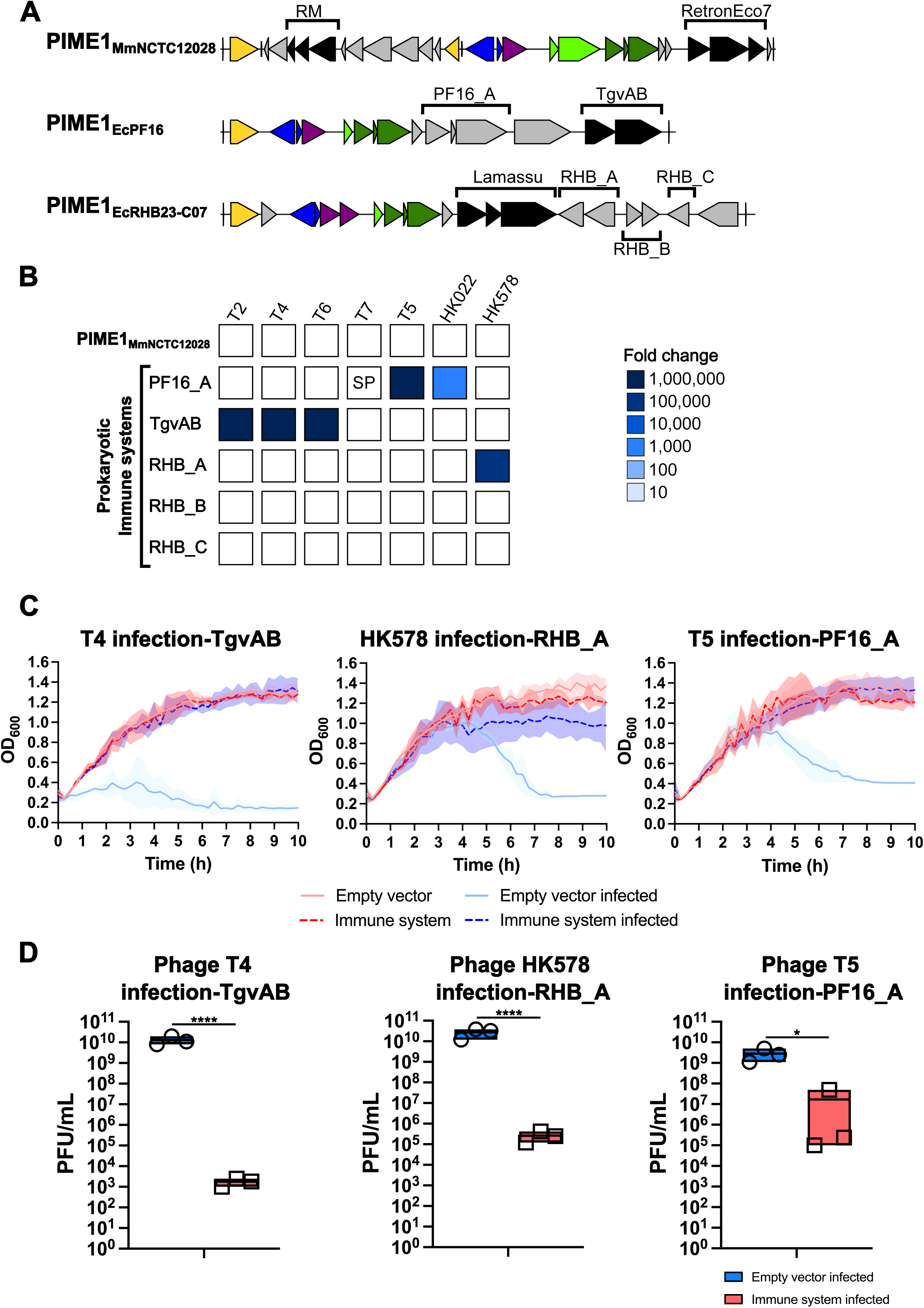
PIMEs are biobanks of multiple prokaryotic immune systems. **(A)** Genomic view of hotspots encoded on PIMEs. Genes are color-coded according to their sequence and predicted function: integrase (*int*) in yellow; transcriptional regulators (*alpA* and *luxR*) in blue; replication genes (*repA* and *repC*) in purple; mobilization genes (*traJ,* and *traI*) in light green; transfer genes (*trbJ*, *trbK,* and *trbL*) in dark green; immune systems in black; and genes encoding hypothetical proteins in grey. Known and putative prokaryotic immune systems tested are highlighted. **(B)** PIME1_MmNCTC12028_, known and putative PIME-immune systems were tested against *E. coli* phages. Heatmap represents the fold-change in phage protection, which was measured using a serial dilution spot plaque assay, comparing the efficiency of each phage to form plaque on strains carrying either the empty plasmid or the plasmid expressing the immune system under study. Data are representative of three replicates. SP represents a small plaque phenotype. **(C)** Strains with empty plasmid or encoding the different immune versions were infected with T4, T5 or HK578 phage at an input MOI (iMOI) of 0.1. **(D)** Phage titers were determined from (C) using *E. coli* C600 or c1a as the recipient. For (D), data were log_10_ transformed prior to analysis. An unpaired two-sided *t* test was performed to compare mean differences between the phage infection vs the empty plasmid and the immune system. Adjusted *p* values were as follows: *ns* > 0.05; \**p* ≤ 0.05; \*\**p* ≤ 0.01; \*\*\**p* ≤ 0.001; \*\*\*\**p* ≤ 0.0001.

Thus, to assess the protective role of PIMEs, we focused on PIMEs present in *E. coli*, concretely PIME1EcRHB23-C07 and PIME1EcPF16, which we have in our strain collection. Instead of using the complete PIME element, we selected known immune systems and adjacent hypothetical regions and cloned them under the control of their native promoters into the pBAD18-kmR plasmid (Fig. 5A). These constructs were introduced into *E. coli* c1a or C600 and tested for anti-phage activity against a diverse panel of seven coliphages spanning various phage families (Table S4). Compared to the empty vector control, three systems conferred robust and reproducible resistance against certain phages (Fig. 5B). Also, recipient strains were infected at an input multiplicity of infection (iMOI) of 0.1 in liquid media and cell survival was measured after phage infection. While strains containing the empty plasmid completely lysed after infection, strains expressing the different immune systems did not reveal significant lysis (Fig. 5C). Then, we quantified the number of infecting particles produced after infection and the phage progeny was severely reduced (Fig. 5D). Importantly, two of the systems exhibited a narrow spectrum, targeting exclusively individual phages or closely related phages, such as T2, T4, and T6 phages for the previously characterized immune system^42,43^, and HK578 for the newly immune system identified RHB_A. In contrast, the newly identified immune system PF16_A displayed a broad defense spectrum, providing protection against unrelated phages (Fig. 5). Collectively, these results demonstrate the idea that PIMEs represent an important reservoir of immune systems and are key contributors to bacterial immunity.

## Discussion

Over the past few decades, our understanding of how non-autonomous MGEs exploit their autonomous counterparts for mobilization has advanced significantly^10,11^. These non-autonomous MGEs have adapted to the two main mechanisms of HGT mediated by MGEs: phages mobilize phage satellites^44–49^, while conjugative elements disseminate mobilizable plasmids and IMEs^12,15,50^. The interplay and conflicts between autonomous and non-autonomous elements and their impact on both bacterial and MGE evolution are key areas of research.

Our study reveals an overlooked and widely disseminated family of IMEs, named PIME (<u>P</u>seudomonadota Integrative <u>M</u>obilizable <u>E</u>lement). This family is composed of several distinct subfamilies that share an *oriT* resembling that of their helper conjugative plasmids and a conserved set of core components involved in integration, regulation, replication, and transfer (Fig. 1 and Fig. S8). Despite this shared core, each subfamily displays a discrete genetic organization and hallmark features, such as the presence of a *luxR* gene in the regulatory region of PIME1 members compared to PIME2. We were able to systematically identify hundreds of PIME elements distributed among diverse Pseudomonadota bacterial species, highlighting their relevance in nature. Notably, PIMEs were detected in environmentally relevant aquatic pathogens such as *Vibrio*, soil-associated opportunistic bacteria including *Citrobacter* and *Acinetobacter*, as well as plant pathogens such as *Xanthomonas*. Yet, our large-scale analysis relies on the co-occurrence of all these core components. However, other PIME variants could exist in nature, lacking one of the defined core components or containing a non-homologous analog component while still being functional.

The origin of PIMEs or how they have evolved remains unclear. Notably, members of different PIME subfamilies can utilize the same *attB* site located adjacent to the *guaA* gene for chromosomal integration (Fig. 1). This feature is not unique to PIMEs, as it also occurs with other MGEs, such as different families of phage satellites^44,49^. However, this observation leads us to speculate that these different PIME subfamilies originated from a common ancestral element, which subsequently underwent genetic reorganizations and acquired specific genes, giving rise to the distinct PIME subfamilies observed. Additionally, conserved homologs of the PIME *repA* gene are also found in mobilizable plasmids, concretely the broad-host RSF1010 plasmid (Fig. S10), which exploits for mobilization the same conjugative plasmids as the PIMEs characterized^51,52^. Because IMEs reside stably integrated in the chromosome while mobilizable plasmids remain extrachromosomal, an important question arises: Could mobilizable plasmids and IMEs that employ the same helper conjugative element share a common origin and have diverged to adapt to distinct lifecycle? What are the advantages to remain integrated or extrachromosomal until the arrival of a helper conjugative plasmid?

Here, we characterized the life cycle of some PIME members belonging to the PIME1 subfamily. Our results indicate that the PIME *int* and regulatory genes are essential for their lifecycle. Indeed, our model system PIME1MmNCTC12028 fails to replicate in the absence of excision and circularization, suggesting the existence of a tight genetic regulation to allow replication and conjugation. This behavior is consistent with previous observations for SGI1, in which replication occurs only after excision and circularization^18,23^. We also observed that helper IncP conjugative plasmids can promote induction of the PIME1EcRHB23-C07 element but do not mediate its mobilization. This lack of mobilization is likely due to the inability of the conjugative plasmid relaxosome to recognize the PIME1EcRHB23-C07 *oriT* region. However, because only two helper IncP conjugative plasmids were tested, we cannot exclude the possibility that other conjugative plasmids from the same IncP group are able to both induce and mobilize this PIME. A similar phenomenon was recently described for phage satellites, in which helper phages promote phage satellite induction but not their transfer, with phage satellites nevertheless being fully assembled extracellularly^53^. Further work is required to determine how they sense the helper conjugative plasmid and whether other PIME subfamilies exploit the same conjugative plasmids and exhibit life cycles similar to that of PIME1.

Additionally, in our model system PIME1MmNCTC12028, we did not observe a reduction in the cotransfer of the helper conjugative plasmid and the PIME element. This observation is contrary to the SGI1 and derivative systems^10,15,54^. However, we observed a reduction in helper conjugative plasmid amplification, suggesting the existence of a potential incompatibility that may be partially compensated. Future work will be needed to determine whether PIME1MmNCTC12028 affects other conjugative plasmids, how mechanistically achieved, and whether conjugative plasmids can deploy compensatory strategies to overcome this effect. This is not the unique layer of interaction between the helper conjugative plasmid and the PIME1MmNCTC12028 element. We also observed that, in contrast to other systems, the helper conjugative plasmid can utilize the relaxase (*traI*) encoded by PIME1MmNCTC12028 for its own mobilization. Conversely, PIME1MmNCTC12028 fails to use the conjugative plasmid encoded relaxase in the absence of its own. Given that some IMEs interfere with plasmid relaxases^25^, this strategy would prevent such negative effect and still allow efficient conjugative plasmid transfer. Whether this mechanism is restricted to the PIME family or is more broadly employed by other IMEs remains to be determined.

Contrary to other non-autonomous MGEs, such as phage satellites, PIMEs are mobilized by conjugation and there is, in principle, no limitation on their size. Nevertheless, most PIME elements have a size of ∼20 kb, and this is consistent with other IME families^15,34,55–57^. This raises the question of why IMEs appear to maintain a constrained genome size. Given recent evidence that phage and phage satellites can shape plasmid evolution^58^, it is tempting to speculate that PIMEs may have retained a size similar to that of phage satellites to enable occasional transfer by transduction.

A striking feature of PIMEs is their large accessory regions, which account for approximately half of their genomes and serve as biobanks of both known and novel prokaryotic immune systems. We further demonstrated that PIMEs can protect their bacterial hosts from phage predation. These observations align with recent studies highlighting MGEs, such as phages and phage satellites, as major contributors to the bacterial defensome^3,59–62^. However, such MGEs exhibit a narrow host range. In contrast, PIMEs are not restricted to specific bacterial hosts and are distributed across phylogenetically distant taxa. This broad distribution is consistent with their mechanism of HGT, as conjugation have a broad host spectrum^63,64^. Thus, the carriage of these genes likely enables the rapid and efficient dissemination of defence systems among bacterial populations and suggest that PIMEs may play a crucial role in the dissemination of prokaryotic immune systems among bacterial populations.

Elucidating how genomic islands mobilize is critical to understand bacterial evolution. However, much of our current knowledge relies on bioinformatic predictions, with relatively few non-autonomous MGEs characterized experimentally. Here, we reveal and characterize the lifecycle of a widespread family of IMEs capable of both intra- and inter-species mobilization. By revealing previously unrecognized layers of interaction between IMEs and their helper conjugative elements, our work improves our understanding of how MGEs interact. We anticipate that future research on hijacker and helper element relationships will uncover additional interaction strategies, shedding light on how these processes shape bacterial and MGEs evolution.

## Material and methods

### Bacterial strains and growth conditions

Conjugative plasmids, phages and bacterial strains used in this study are listed in Table S3, S4 and S5, respectively. *E. coli* and *M. morganii* strains were grown at 30°C or 37°C on Luria-Bertani (LB) agar plates or in LB broth with shaking (120 rpm). When required, media were supplemented with ampicillin (100 µg mL^−1^), kanamycin (30 µg mL^−1^), chloramphenicol (*cat*) (20 µg mL^−1^), or tetracycline (*tetA*) (20 µg mL^−1^), form Sigma-Aldrich.

### Plasmid construction

Plasmids generated in this study, listed in Table S6, were constructed by cloning PCR-amplified products, obtained using the oligonucleotides listed in Table S7, into the corresponding vectors (pBAD18, pBAD18-*kmR*, and pK03Blue) using standard restriction enzyme digestion and ligation procedures. Plasmid constructs were verified by Sanger sequencing in Eurofins Genomics.

### DNA methods

For *E. coli* and *M. morganii*, gene insertions and deletions were performed as previously described using λ Red recombinase-mediated recombination^44,65,66^. Chloramphenicol (*cat*), tetracycline (*tetA*), or kanamycin (*kmR*) resistance markers were inserted into the bacterial chromosome, IMEs or conjugative plasmids. Briefly, resistance markers were PCR-amplified using primers listed in Table S7 from plasmids pKD3, pKD4, or from a strain carrying the *tetA* marker. Then, PCR products were transformed into recipient strains harboring plasmids pRWG99 or pRWG99-*kmR*, which express the λ Red recombinase, facilitating markers integration into the bacterial chromosome, IME, or conjugative plasmid genomes. When excision of resistance markers from the IME or conjugative plasmid was required to generate scarless deletions, plasmid pCP20 was transformed into the corresponding strains. Cultures carrying pCP20 were grown overnight at 30°C, followed by a 1:50 dilution into fresh LB broth, and incubated at 42°C for 4 h to encourage plasmid loss, while allowing FLP-mediated recombination. Cells were then plated out on LBA plates and incubated at 37°C. Single colonies were streaked out, and PCR was used to corroborate excision of the resistance marker. The different mutants obtained were verified by PCR and Sanger sequencing in Eurofins Genomics.

Scarless deletion of IMEs was performed using site-directed scarless mutagenesis as described previously^67,68^. Briefly, the *kmR* or *cat* resistance marker, together with an I-*Sce*I recognition restriction site, was PCR-amplified using primers listed in Table S7. Then, the PCR product was introduced into the recipient strain harboring plasmid pRWG99 or pRWG99-*kmR*, which express the λ Red recombinase. Following successful recombination, 300bp PCR fragments amplified from the empty *attB* site (*guaA* gene) were electroporated into the mutant strain expressing the λ Red recombinase. Successful recombinants were selected based on the expression of I-*Sce*I endonuclease. Successful IME deletion mutants were confirmed by PCR and Sanger sequencing in Eurofins Genomics.

### Conjugation and mobilizable plasmids assay

To measure transconjugation efficiency, mating assays were performed by mixing equal volumes of recipient and donor strains. Recipient strains carried the appropriate antibiotic resistance marker, while donor strains harbored the corresponding conjugative plasmid along with either an IME (and their plasmid complementation) or a mobilizable plasmid. Overnight cultures of donor and recipient strains were diluted 1:50 in fresh LB broth and grown to an OD600 = 1. For complementation experiments, 0.02% arabinose was added to the medium to induce overexpression of the gene of interest. Then, cells were harvested by centrifugation to remove antibiotics from the medium and resuspended in 1 mL of fresh LB broth. For the mating assays, 100 µL of the recipient strain was mixed with 100 µL of the donor strain and spotted onto LB agar plates. Plates were incubated at 37°C for 3 h to allow conjugation. Following incubation, cells were resuspended in 1 mL of LB broth, and serial dilutions were prepared and plated out onto LBA plates containing the appropriate antibiotics to quantify donor, recipient, and transconjugants corresponding to the conjugative plasmid, IME, or mobilizable plasmid. Transconjugation frequency was determined as the number of transconjugants per recipient cells (TR/R) in the mating mixture at the time of plating.

### qPCR

Genomic DNA was extracted from 1.5 mL of cell culture using the GenElute^TM^ Bacterial Genomic DNA Kit from Sigma-Aldrich. DNA purity and concentration were determined using an One^C^ NanoDrop spectrophotometer (Thermo Fisher Sientific). qPCR experiments were performed in biological triplicates using the Power Up SYBR Green Master mix (Thermo Fisher Sientific) on an AriaMx Real-Time PCR system (Agilent Technologies). Reactions were performed in a total volume of 10 μL in 96-well plates, containing 5 μL of master mix, 3μL of DNA (6 ng), and 2 μL of primer pair solutions (1 μM). Primers used in qPCR analyses are listed in Table S7. Cycling conditions were as follows: 3 min at 95°C; 50 cycles: 15 s at 95°C, 30 s at 60°C, and 40 s at 72°C; followed by a final elongation step of 3 min at 72°C. Data were analyzed using the 2^-ΔCT^ method (where CT is the threshold cycle) with *hofN* gene as the chromosomal reference.

### Phage plaque assay

*E. coli* strains carrying either the empty pBAD18-*kmR* plasmid or pBAD18-*kmR* encoding different immune systems were grown overnight. Overnight cultures were diluted 1:50 in fresh LB broth and grown to an OD600 = 0.34. Bacterial lawns were prepared by mixing 100 µL of cells with phage top agar (PTA; 20 g Nutrient Broth No. 2, Oxoid; 3.5 g agar, 10 mM MgCl2) and poured onto phage base agar plates (PBA; 25 g of Nutrient Broth No. 2, Oxoid; 7g agar) supplemented with 0.02% arabinose and 10mM CaCl2. Serial dilutions of phages were prepared in phage buffer (50mM Tris pH 8, 1mM MgSO4, 4mM CaCl2 and 100mM NaCl) and spotted onto the corresponding bacterial lawns. Plates were incubated at 37°C for 24 h. The fold change in protection was calculated by quantifying the number of plaques formed on the strain carrying the empty plasmid divided by the number of plaques on the strain carrying the immune system. The titration limit of detection is 10 plaque forming units (PFUs). SP means small plaques.

### Growth curves assay

*E. coli* strains carrying either the empty pBAD18-*kmR* plasmid or pBAD18-*kmR* encoding different immune systems were grown overnight. Overnight cultures were diluted 1:50 in fresh LB broth and grown to an OD600 = 0.34. Phage was added at an iMOI of 0.1, and 0.02% of arabinose was added to induce expression from the pBAD18 plasmid. Infected cultures were seeded into 96-well clear bottom plates (Corning) and incubated in a Varioskan Lux fluorometer plate reader (Thermofiher) at 37°C with shaking at 500 rpm. OD600 was measured every 15 min to monitor bacterial growth.

### Phage titration

Phage lysates obtained from infections were filtered using sterile 0.2 μm filters (Minisart® single use syringe filter unit, hydrophilic and non-pryogenic, Sartonium Stedim Biotech). The number of phage particles was quantified using the tittering assay. *E. coli* strains were grown overnight. Overnight cultures were diluted 1:50 in fresh LB broth and grown to an OD600 = 0.34. Then, 50 µL of cells were mixed with 100 µL of phage lysate diluted in phage buffer and incubated at room temperature for 10 min. Subsequently, 3 mL of PTA was added, and the infection mixture was poured out onto PBA plates supplemented with 5mM of CaCl2. Plates were incubated at 37°C for 24 h. The number of plaques formed, indicative of phage particles present in the lysate, were counted and represented as PFU/mL. The titration limit of detection is 10 PFUs.

### Identification of novel IMEs and phylogenetic analysis

We retrieved the latest release of the NCBI RefSeq database (version 232; December 1^st^, 2025), comprising a total of 446,233 bacterial genomes. Complete and draft genomes were employed to identify putative IMEs, while plasmids and low-quality contigs were discarded from the analysis. Hidden-Markov model (HMM)^69^ profiles were created for core components (*int, alpA, repA*, *trbJ*, and *trbL*) and used to search for homologues across the RefSeq database. Only positive hits with e-values < 1e-5 were retained. Custom scripts (available at https://github.com/francisconadalm/GenomeRadar) were used to obtain information on bacterial taxa and genome features, including genomic position, locus, accession numbers, gene products, and taxonomic assignment, as well as to identify co-occurrence of core components. Only putative IMEs containing a complete set of core components identified were retained for further analysis (Table S2). Core proteins were individually aligned to their homologs using MAFFT (v7.505)^70,71^, and the resulting alignments were concatenated using a custom script (https://github.com/francisconadalm/GenomeRadar). A phylogenomic tree was constructed using IQ-TREE2 (v2.0.7) with partitioned model selection under the parameters “-m MFP -bb 10000 -alrt 1000”^72,73^. The final model implemented by IQ-TREE2 was VT+F+R7. The resulting phylogenomic tree was visualized and annotated using iTOL^74^.

To classify a PIME element into a specific subfamily, each putative element was aligned with the integrase positioned at the left end. The following criteria were applied: (i) PIME1 and PIME2 subfamilies: elements with core components in the same orientation as the integrase were assigned to either PIME1 or PIME2. PIME1 elements were further distinguished by the presence of a *luxR* gene in the regulatory region; (ii) PIME3 subfamily: elements with regulatory (*alpA*) and replicase (*repA*) genes in the same orientation as the integrase, and transfer genes (*trbJ* and *trbL*) in the opposite orientation, were classified as PIME3; and (iii) PIME4 subfamily: elements with core components in the opposite orientation to the integrase were classified as PIME4. These elements also encode the regulatory (*alpA*) and replicase (*repA*) genes at the right end of the element.

### Whole genome sequencing (WGS) and analysis

Genomic DNA was extracted from 1.5 mL of cell culture using the GenElute^TM^ Bacterial Genomic DNA Kit from Sigma-Aldrich. DNA quality and integrity were tested using an Agilent Bioanalyzer 2100. DNA libraries were prepared using the Nextera XT sample preparation kit, and WGS was performed at the FISABIO Sequencing Facility using the Illumina NextSeq platform (San Diego, CA, USA) to generate 2 x 150 bp paired end reads. Only one replicate was sequenced per experiment.

The quality of sequencing reads was initially assessed using FastQC v0.11.9^75^ and MultiQC v1.11^76^. Adapter sequences and low-quality reads were removed using PrinSEQ v0.20.4^77^. Sequencing reads were then mapped to their respective reference genomes using the Burrows-Wheeler Alignment (BWA) v0.7.17^78^: *E. coli* C600 (CP031214), pB10 (AJ564903), and PIME1MmNCTC12028 (LS483498; positions 1.652.603-1.675.945). BAM files were generated with Picard-tools v2.1.1, merged using SAMtools v1.11^79^, and subsequently sorted and indexed. Bed files were produced using BEDtools v2.30.0 with the *bamtobed* subcommand^80^. Relative coverage across the chromosome was calculated using 100 bp sliding windows. For each experiment, the average coverage of the full genome (excluding pB10 and PIME1MmNCTC12028 sequences, removed with bedtools subcommand *subtract*) was computed. Coverages within each sliding windows was then divided by the chromosomal average to obtain the relative coverage. Reads mapping to the chromosome, pB10, and PIME1MmNCTC12028 were quantified and represented as depth of coverage relative to the average coverage of the entire genome.

### Quantification and statistical analysis

For all quantitative data, experiments were repeated at least three times, with sample sizes indicated in the figure legends. Data are presented as mean ± standard deviation (SD). Statistical analyses were performed as indicated in the figure legends using GraphPad Prism v10.4.1 software. One-way ANOVA with Dunnett’s multiple comparisons test was applied to compare three or more groups. An unpaired t-test was performed to compare two groups. Adjusted p values as: *ns* > 0.05; ∗*p* ≤ 0.05; ∗∗*p* ≤ 0.01; ∗∗∗*p* ≤ 0.001; ∗∗∗∗*p* ≤ 0.0001.

## Data availability

The WGS data generated in this study have been deposited in the NCBI SRA database under accession code BioProject.

## Acknowledgments

We thank José R. Penadés and Lingchen He (Imperial College London, UK) for his support and advice on this project. We thank Nicole Stoesser for sharing bacterial strains (University of Oxford, UK). This work was supported by a PID2023-152560NA-I00 from Spanish Government (Ministry of Science and Innovation) and a CDEIGENT Research Fellowship from Valencian Government [CIDEIG/2023/18] to A.F.-S. A.M.P. was supported by an FPI predoctoral fellowship from the Spanish Ministry of Science and Innovation.

## Author contributions

A.F.-S. conceived the study; A.M.P. and A.F.-S. conducted the experiments; F.N.-M. and P.J.C.-Y. performed the genomic analyses; A.M.P., F.N.-M., P.J.C.-Y., F.G.-C. and A.F.-S analyzed the data. A.F.-S. wrote the manuscript with input from other authors.

